# Alternate without alternative: Neither preference nor simple learning behaviour shown by C57BL/6J mice in the T-maze

**DOI:** 10.1101/2020.11.11.377788

**Authors:** Anne Habedank, Pia Kahnau, Lars Lewejohann

## Abstract

In rodents, the T-maze test is commonly used to investigate spontaneous alternating behaviour but it can also be used to investigate memory, stimuli discrimination or preference between goods. However, especially regarding T-maze preference tests there is no recommended protocol and researchers frequently report reproduction difficulties of this test using mice.

Here, we aimed to develop an efficient protocol with female C57BL/6J mice, conducting two preference tests with different design: In a first test, on two consecutive days with five trials, thirteen mice had to choose between two fluids. In a second preference test, on five consecutive days with two (week 1) or three (week 2) trials, twelve mice had to choose between one arm containing bedding mixed with millet and one containing only bedding. This test design resembled a simple learning test (learn where to find the rewarded and the unrewarded arm on the basis of spatial, olfactory and visual cues).

In both experiments, mice took only a few seconds per trial to run the maze and make their choice. However, in both experiments mice failed to show any preference for one of the arms. Instead, they alternated choices. We therefore believe the T-maze test to be rather unsuitable to test preference or learning behaviour with C57BL/6J mice.

## 1 Introduction

The T-maze is a behavioural test using a maze with a start arm (sometimes connected to a start cage) and two choice arms branching of at the same point from the start arm. In the classic design the arms lie exactly opposite each other, so that they form a T together with the starting arm. In the Y-maze variation, the arms branch off from the start arm at a steeper angle so that the overall shape of the apparatus is y-shaped. During a T-maze test, an animal is placed either in the start cage or directly inside the maze at the beginning of the start arm. At the end of the start arm, the animal has then to choose between entering the left or the right arm. Depending on the setup, in addition to the spatial position the arms can provide further cues, e.g., visual (mice: Lione et al. 1999; broilers: Buckley et al. 2011), tactile (compare Cunningham et al. 2006) or olfactory cues (Mayeux-Portas et al. 2000). Also, none, one or both arms can contain a reward, which can be food (Deacon and Rawlins 2006; Deacon 2006; Crusio et al. 1990), shelter (Pilz et al. 2020) or a platform (in case of the water T-maze, Guariglia and Chadman 2013; Granholm et al. 2000; Belzung et al. 2001).

The T-maze is an important behavioural test to assess the effect of drugs (mice: Correa et al. 2015; rats: Lohninger et al. 2001), genetic alterations (mice: Mayeux-Portas et al. 2000; Granholm et al. 2000) or diseases (mice: Belzung et al. 2001; rats: Sánchez-Santed et al. 1997; Wu et al. 2018). It is often used to assess spontaneous alternating behaviour, spatial memory and/or discrimination of stimuli (Deacon and Rawlins 2006; Deacon 2006; Dudchenko 2004; Dember and Fowler 1958; Sharma et al. 2010b; Wenk 1998). Spontaneous alternating behaviour describes the tendency of rodents to choose the arm they did not visit in the preceding trial. This kind of behaviour occurs spontaneously and is not necessarily related to a resource being exploited in the preceding trial (mice: Gerlai 1998; gerbils: Dember and Kleinman 1973; rats: Sánchez-Santed et al. 1997). In position discrimination tests (also: spatial memory tests), only one spatial location, either the left or the right arm, is baited (mice: Guariglia and Chadman 2013; Lione et al. 1999; Pioli et al. 2014; Sharma et al. 2010a; Granholm et al. 2000; Belzung et al. 2001). Thus, the spontaneous alternating is a way to evaluate the short-time or working memory (which location was last visited?), while the position discrimination test evaluates the reference memory, similar to the conditioned place preference test (Sharma et al. 2010b; Shoji et al. 2012; Wenk 1998; Hieu et al. 2020). In a further modification of the position discrimination, the T-maze can also be used as general discrimination test, using additional cues instead of merely the spatial one to provide information on the baited arm (mice: Lione et al. 1999; Mayeux-Portas et al. 2000; Granholm et al. 2000; broilers: Buckley et al. 2011).

In a modification of the discrimination test, the T-maze can also be used as a preference test: The arms are provided with different goods, and the animal is required to choose between them. This form of preference test seems to be easily performed with a variety of animal species (mice: Cutuli et al. 2015; Correa et al. 2015; Roder et al. 1996; wild mice: Nunes et al. 2009; rats: Leenaars et al. 2019; Ras et al. 2002; Patterson-Kane et al. 2001; van der Plasse et al. 2007; Denk et al. 2004; Hernandez-Lallement et al. 2015; Wadhera et al. 2017; Cunningham et al. 2015; pigs: Rooijen and Metz 1987; hens: Dawkins 1977; broilers: Buckley et al. 2011; zebrafish: Hieu et al. 2020; fruit flies: Fujita and Tanimura 2011). Preference is usually assessed by offering the goods in the choice arms of the maze but in some cases, it might be useful to use stimuli which are associated with the to-be-tested goods instead, e.g., in tests for social preference, the real mouse might be replaced by urinary stimuli (Nunes et al. 2009; compare also Fitchett et al. 2006). It also has to be kept in mind that offering the goods itself can lead to saturation and/or influence the choice in the next trial (Kirkden and Pajor 2006), in the same way as humans might prefer milk after eating something spicy (Nasrawi and Pangborn 1990).

Preference tests in T-mazes can be performed with discrete or continuous choices: In a discrete measurement, an animal has to perform multiple trials in which it can choose between the left or the right arm (mice: Tellegen et al. 1969; rats: Patterson-Kane et al. 2001; Pioli et al. 2014; Ras et al. 2002; van der Plasse et al. 2007). In a continuous measurement, the animal stays in the T-maze for a defined period of time and the time the animal spends in the left or the right arm is used to ascertain preference (mice: Cutuli et al. 2015; Roder et al. 1996; wild mice: Nunes et al. 2009; Correa et al. 2015; compare also Pennycuik and Cowan 1990; using a U-shaped maze and wild mice).

There are various protocols and recommendations on the conduction of T-maze tests for behavioural measures such as memory and discrimination. However, there is to date no protocol for T-maze preference tests: The protocols focus either on spontaneous (unrewarded) alternation (Deacon and Rawlins 2006; Wenk 1998), rewarded alternation (Deacon and Rawlins 2006; Shoji et al. 2012; Wenk 1998) or position discrimination (Deacon 2006; Shoji et al. 2012). A short comparison of different protocols is given in Table 1. In general, for spontaneous alternation, no food restriction or habituation is needed. Animals should just be well-habituated to their environment and the handling, before they are placed into the maze. Protocols for rewarded alternation and position discrimination are more complex and differ in their recommendations. Often, food restriction to 85 % of free-feeding weight is recommended, although Deacon and Rawlins (2006) at the same time states that well habituated animals should also perform the T-maze without food restriction (Deacon and Rawlins 2006). For rewarded alternation, forced trials are recommended, in which animal are only allowed to visit one arm by blocking the other. In the following trial, animals get a free choice with both arms accessible. If the animals visit the previously blocked arm, they made an alternating choice. In position discrimination on the other hand, no forced trials are conducted, and trials are always free choice. Also, rewarded alternation and position discrimination differ with regard to the recommendations made about cleaning: While in for rewarded alternation tasks, cleaning seems to be more common, for position discrimination Deacon (2006) explicitly states that no cleaning maximizes the learning potential (Deacon 2006). However, protocols for both types of tests differ greatly in their recommendations for habituation procedure (individuals or group, duration, free exploration or trials, reward or no reward) and intertrial interval (immediately or more than 10 min). All protocols recommend at least ten trials per day, but depending on the intertrial interval this leads to differing test durations from 50 min (Shoji et al. 2012) to several hours (Deacon 2006). None of the protocols gives instructions with regard to testing time, although day time might influence motivation to gain food (Acosta et al. 2020; Koch et al. 2020).

**Table 1:**
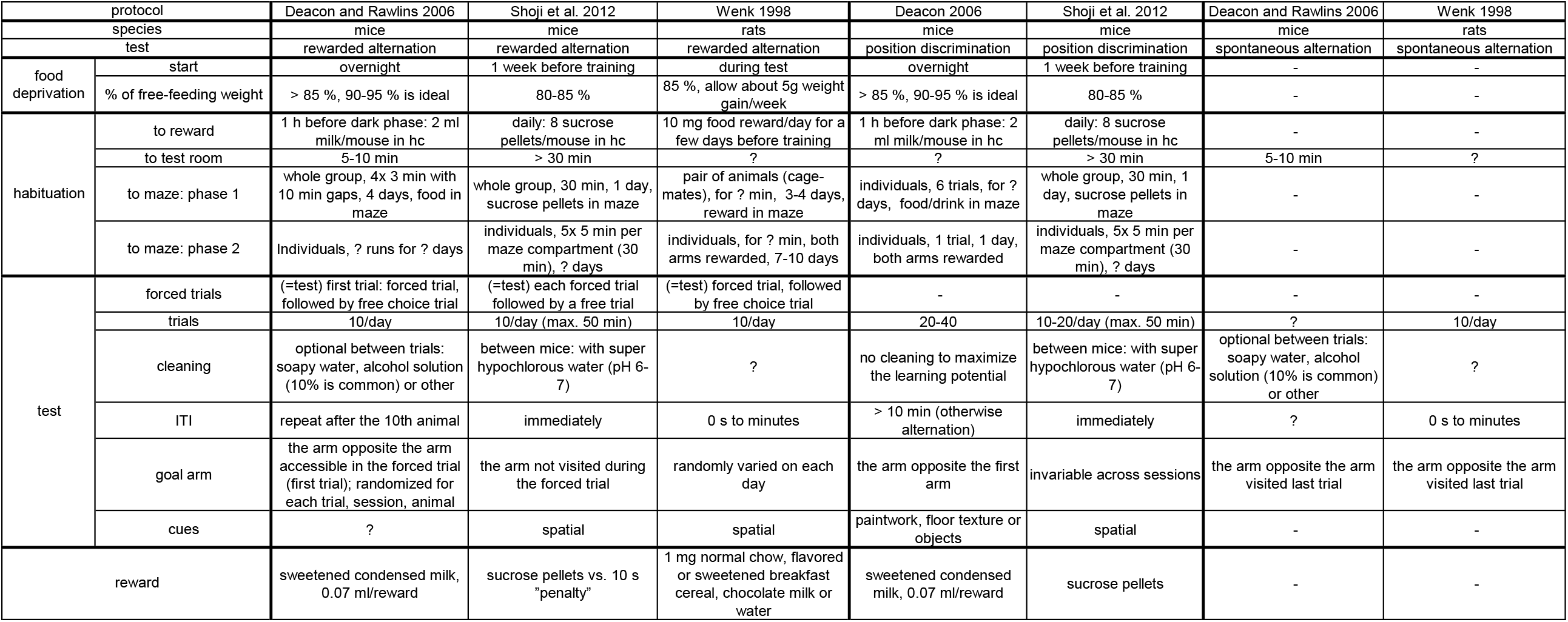
Comparison of T-maze protocols by Deacon and Rawlins 2006, Deacon 2006, Shoji et al. 2012 and Wenk 1998. ITI = intertrial interval, hc = home cage, ? = not described see extra table file “T-Maze_Protocols.pdf”

| protocol |  | Deacon and Rawlins 2006 | Shoji et al. 2012 | Wenk 1998 | Deacon 2006 | Shoji et al. 2012 | Deacon and Rawlins 2006 | Wenk 1998 |
| --- | --- | --- | --- | --- | --- | --- | --- | --- |
| species |  | mice | mice | rats | mice | mice | mice | rats |
| test |  | rewarded alternation | rewarded alternation | rewarded alternation | position discrimination | position discrimination | spontaneous alternation | spontaneous alternation |
| food deprivation | start | overnight | 1 week before training | during test | overnight | 1 week before training | - | - |
|  | % of free-feeding weight | > 85 %, 90-95 % is ideal | 80-85 % | 85 %, allow about 5g weight gain/week | > 85 %, 90-95 % is ideal | 80-85 % | - | - |
| habituation | to reward | 1 h before dark phase: 2 ml milk/mouse in hc | daily: 8 sucrose pellets/mouse in hc | 10 mg food reward/day for a few days before training | 1 h before dark phase: 2 ml milk/mouse in hc | daily: 8 sucrose pellets/mouse in hc | - | - |
|  | to test room | 5-10 min | > 30 min | ? | ? | > 30 min | 5-10 min | ? |
|  | to maze: phase 1 | whole group, 4x 3 min with 10 min gaps, 4 days, food in maze | whole group, 30 min, 1 day, sucrose pellets in maze | pair of animals (cage-mates), for ? min, 3-4 days, reward in maze | individuals, 6 trials, for ? days, food/drink in maze | whole group, 30 min, 1 day, sucrose pellets in maze | - | - |
|  | to maze: phase 2 | Individuals, ? runs for ? days | individuals, 5x 5 min per maze compartment (30 min), ? days | individuals, for ? min, both arms rewarded, 7-10 days | individuals, 1 trial, 1 day, both arms rewarded | individuals, 5x 5 min per maze compartment (30 min), ? days | - | - |
| test | forced trials | (=test) first trial: forced trial, followed by free choice trial | (=test) each forced trial followed by a free trial | (=test) forced trial, followed by free choice trial | - | - | - | - |
|  | trials | 10/day | 10/day (max. 50 min) | 10/day | 20-40 | 10-20/day (max. 50 min) | ? | 10/day |
|  | cleaning | optional between trials: soapy water, alcohol solution (10% is common) or other | between mice: with super hypochlorous water (pH 6-7) | ? | no cleaning to maximize the learning potential | between mice: with super hypochlorous water (pH 6-7) | optional between trials: soapy water, alcohol solution (10% is common) or other | ? |
|  | ITI | repeat after the 10th animal | immediately | 0 s to minutes | > 10 min (otherwise alternation) | immediately | ? | 0 s to minutes |
|  | goal arm | the arm opposite the arm accessible in the forced trial (first trial); randomized for each trial, session, animal | the arm not visited during the forced trial | randomly varied on each day | the arm opposite the first arm | invariable across sessions | the arm opposite the arm visited last trial | the arm opposite the arm visited last trial |
|  | cues | ? | spatial | spatial | paintwork, floor texture or objects | spatial | - | - |
| reward |  | sweetened condensed milk, 0.07 ml/reward | sucrose pellets vs. 10 s "penalty" | 1 mg normal chow, flavored or sweetened breakfast cereal, chocolate milk or water | sweetened condensed milk, 0.07 ml/reward | sucrose pellets | - | - |

Thus, with regard to rewarded alternation and position discrimination, there is not one “perfect” test design. Moreover, personal correspondence with other researchers resulted mainly in reports of difficulties in reproduction of the T-maze test, even more so, when researchers tried to alter the existing protocols for preference tests. In general, varying success rates might be caused by differences in strain performances (Gerlai 1998; Moy et al. 2008). However, there are various additional factors which might influence results, e.g., differences in handling technique (base of the tail compared to cup or tube handling, Hurst and West 2010; Gouveia and Hurst 2017), stress (Mitchell et al. 1985), habituation (Rudeck et al. 2020; Deacon and Rawlins 2006), level of food restriction (Richman et al. 1986).

One interesting solution for the factor handling is provided by Zhang et al. (2018), which developed an automated T-maze system (Zhang et al. 2018). Here, no handling is involved, and thus, influence of the researcher is reduced. Taking it one step further, Pioli et al. (2014) introduced an automated T-maze which is even home cage based. Here, mice can conduct the test when active and most motivated to work for the reward, which also makes food restriction superfluous (Pioli et al. 2014). However, this automated T-maze is designed for single housing (there is only a companion animal behind a partition), which might not be the desired husbandry condition. In addition, this automated T-maze is originally designed for spontaneous alternation tasks and it would probably need adjustments for preference tests with regard to, e.g., cue presentation and change of presentation side.

Thus, a working protocol for the conduction of a T-maze preference test is still needed. Here, we performed two experiments in search for such a protocol: In experiment 1, we investigated the preference between two fluids (apple juice vs. almond milk). In experiment 2, we changed the test design and offered one arm containing millet and bedding, and one arm containing only the bedding.

## 2 Material and Methods

### 2.1 Animals

A group of thirteen female C57BL/6J CrL mice was purchased in December 2017 at the age of 3 weeks from Charles River, Sulzfeld. This group was used in experiment 1 (“group 1”). Another group consisting of twelve female C57BL/6J CrL mice was purchased in June 2019 at the age of 4 weeks from Charles River, Sulzfeld. This group was used for experiment 2 (“group 2”). All mice within a group had different mothers and different nurses to ensure maximal behavioural variability within the inbred strain. At the age of five weeks, transponders were implanted, a procedure performed under anesthesia and analgesia (for details see Supplements). Both groups took part in multiple other experiments, including the development of an home cage based automated tracking system and conditioned place preference tests. By the time the T-maze test was performed, they were around 12 months (group 1) or 11 months old (group 2). Both groups were always handled by tube handling. It has to be noted that by the start of the experiment 2, eleven of twelve mice in group 2 at least partly lacked their whiskers. However, this should in theory not have influenced their ability to perceive the visual, olfactory or spatial cues and to act on them.

### 2.2 Housing

One group of mice was kept in two type IV macrolon cages (L × W × H: 598 × 380 × 200 mm, Tecniplast, Italy) with filter tops. The two cages were connected via a Perspex tube (40 mm in diameter). This cage system was chosen because of other research purposes, and mice had lived in it since they were around 2 months (group 1) or 3 months old (group 2). Food (autoclaved pellet diet, LAS QCDiet, Rod 16, Lasvendi, Germany) and tap water (two bottles each cage) were available ad libitum in both cages. Cages were equipped each with bedding material (Lignocel FS14, spruce / fir, 2.5-4 mm, JRS, J. Rettenmaier & S öhne GmbH + Co KG, Germany) of 3-4 cm height, a red house (The Mouse-House, Tecniplast), papers, cotton rolls, strands of additional paper nesting material, and two wooden bars to chew on. Both cages also contained a Perspex tube (40 mm in diameter, 17 cm long), which was used for tube handling. Room temperature was maintained at 22 ± 3 °C, the humidity at 55 ± 15 %. Animals were kept at 12h/12h dark/light cycle with the light phase starting at 7:00 a.m. (winter time) or 8:00 a.m. (summer time), respectively. Between 6:30 and 7:00 a.m. (winter time) or 7:30 and 8:00 (summer time) a sunrise was simulated using a Wake-up light (HF3510, Philips, Germany). Once per week, the home cages were cleaned and all mice were scored and weighed. In this context, mice also received a colour code on the base of their tails, using edding 750 paint markers, to facilitate individual recognition.

### 2.3 T-Maze Setup

For the T-maze test, a start cage (type III, L × W × H: 425 × 266 × 155 mm, Tecniplast, Germany) filled with 1 cm bedding was connected via a tube to the T-maze. The tube contained an automated door. In experiment 1, the connection between the start cage and the T-maze resembled part of the setup used for habituation so mice were already habituated to it (compare Fig. 1a and Fig. 1b): a 15 cm tube with an RFID antenna between cage and door, a 6 cm tube with a light barrier between door and maze. If the mouse interrupted the light barrier in front of the door or was detected by the RFID antenna, the door opened for 5 seconds. For experiment 2 (without automated habituation), the tube connected to the start cage was 14 cm long and contained an RFID antenna, followed by the automated door and a 1 cm long tube (see Fig. 1c). Here, the door also opened for 5 s whenever the transponder of a mouse was detected. There was no light barrier on the other side of the door because this time mice were not allowed to return to the start cage by themselves.

**Fig. 1:**
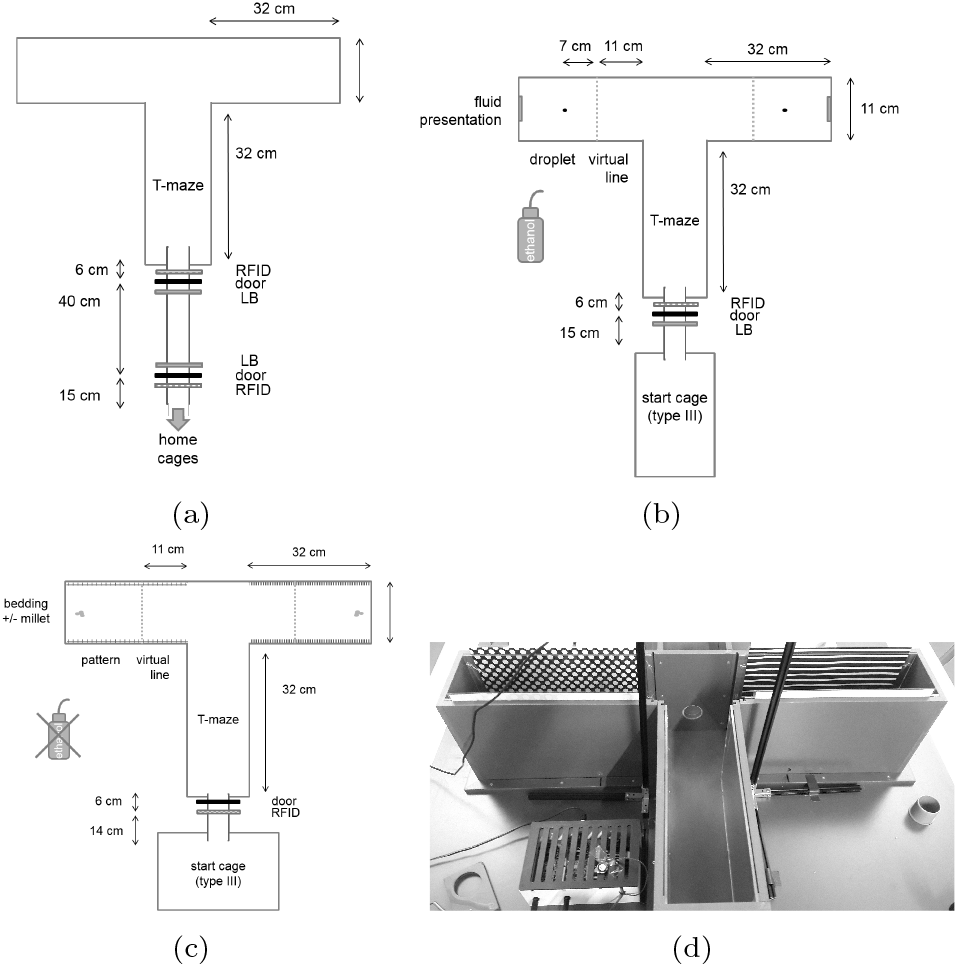
T-Maze setup as a schematic drawing for experiment 1 habituation (a) and test (b), and test of experiment 2 (c). (d) photo of the experiment 2 setup, the box on the bottom left contains the Arduino, which operates the automatic door, the device to its left (with the hole) is an example of the RFID antenna and light barrier constructions. LB: light barrier, door: automatic door, RFID: RFID antenna.

The T-maze itself consisted of gray plastic and had three arms, each 32 cm long and 11 cm wide, with 20 cm high walls (see Fig. 1d). On either side of the arms a mark was made outside the T-maze so that a virtual line could be drawn 11 cm from the central arm during video analysis. If a mouse crossed this line with its whole body (but not yet with its tail), this was defined as a choice being made.

For video recording, in both experiments a webcam (Logitech C390e, Switzerland) was mounted above the maze on a metal beam construction. The connected computer was placed near the T-maze in such a way that the experimenter could observe the mouse in the T-maze via the computer screen.

### 2.4 T-Maze Test

In the first experiment, the T-maze test was used to compare the preference for two fluids. Mice performed discrete choices between the two arms, which contained a droplet of either almond milk or apple juice. Because insufficient habituation might slow the performance in the maze (Deacon and Rawlins 2006) and might be one of the main problems, we conducted a thorough habituation phase: For about two weeks, mice had free access to the T-maze via a connection to the home cage. After one week, fluids were presented for 24 h inside the home cage. After thirteen days, mice were moved to the testing room, to habituate to it before the start of the actual T-maze test. The test was then performed on two days, with five test trials per mouse per day and a side change after the seventh trial to control for side preference (see Fig. 2a). The mice had the choice between almond milk and apple juice, with 20 *µ*l of fluid as a reward in the respective arm. As an olfactory cue, we applied some of the fluid onto a cellulose sheet at the end of the arms. Between mice, the maze was cleaned with ethanol. During trials, an additional light was added. (For more details on the procedure see Supplements.)

**Fig. 2:**
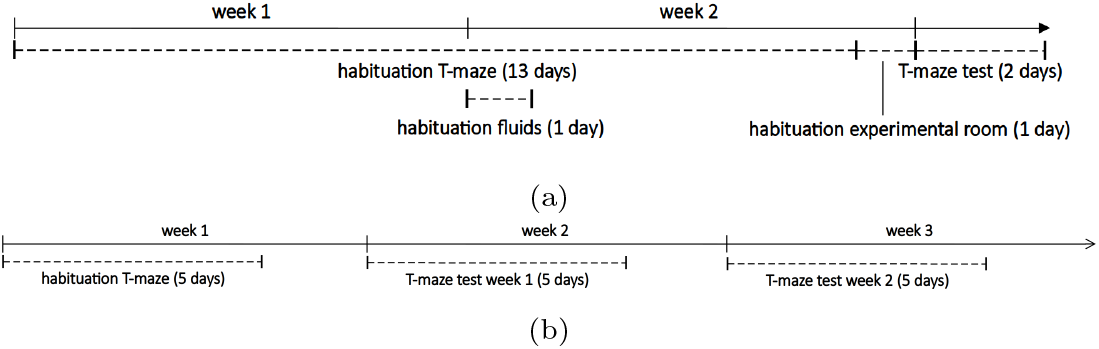
Timeline of experiment 1 (a) and experiment 2 (b). In experiment 2, no habituation to the experimental room was necessary because it took place in the husbandry room. In addition, no habituation to the options (millet with or without bedding material) was necessary because mice were familiar with it from previous experiments.

In a second experiment we changed the design in several points (see Table 2): active (manual) habituation instead of passive habituation for 3 min on five consecutive days, daily repeated trials instead of block-wise trials, no ethanol disinfection of the maze in-between mice, no additional light for the T-maze, and visual cues supplementary to olfactory cues. Also, the choice was now not between two fluids but between millet (0.1 g mixed with bedding material) or no millet (a visually similar amount of bedding material). This preference test therefore resembled a learning test because only one arm was baited. The test was performed on five consecutive days with two trials per mouse per day (leading to the same amount of trials as in experiment 1) and a side change after the sixth trial. Then, after this proved not to show the hoped-for results, a second week was added (see Fig. 2b): Again the test was conducted on five consecutive days but this time three trials were conducted per mouse per day (i.e. one trial more than there were options) and there was no side change (thus, spatial and visual cue provided the same information). A comparison of the timeline of both experiments can be found in Fig. 2. (For more details on the procedure see Supplements.)

**Table 2:** Experimental design of the T-maze tests conducted in experiment 1 and 2. Further explanations on the procedures can be found in the Supplements.

|  |  | Experiment 1 | Experiment 2<br>week 1 | Experiment 2<br>week 2 |
| --- | --- | --- | --- | --- |
| **habituation procedure** | method<br>duration<br>habituation trial | passive<br>13 days<br>1 (on day 1) | active<br>5 days<br>no |  |
| **general test setup** | cleaning<br>illumination<br>options | with 70 % ethanol<br>171-350 lux<br>almond milk<br>vs. apple juice | no<br>18-50 lux<br>millet + bedding<br>vs. bedding |  |
| **test procedure** | duration<br>trials/day/mouse<br>side change<br>cue | 2 days<br>5<br>after trial 7 (day 2)<br>odour | 5 days<br>2<br>after trial 6 (day 4)<br>pattern<br>(+ odour) | 5 days<br>3<br>no<br>pattern + side<br>(+ odour) |

### 2.5 Statistical Analysis

In short, for the T-maze preference test video recordings were analysed with the help of BORIS (Behavioral Observation Research Interactive Software, Version 7.9.8, Friard and Gamba 2016), noting the time points a) when the mouse was placed into the start cage (only experiment 2), b) when the mouse entered the maze, c) when the mouse crossed the virtual line in one of the choice arms, 11 cm into the arm, d) when it entered the handling tube to be returned to the start cage or the home cage. Each behaviour was only counted when the mouse had all four paws on the bedding of the start cage (only experiment 2) or the whole mouse (except the tail) had entered the maze, the tube, or crossed the virtual line (both experiments).

All time points and choices were filled into a table and further managed with the help of R studio (experiment 1: Version 1.1.383, experiment 2: Version 1.2.1335, using R 3.4.0 or higher). For each mouse, choices were pooled (experiment 1: for both days, experiment 2: per week), and the percentage of choices for one option was calculated. Examined were side preference (left vs. right), the option preference (almond milk vs. apple juice in experiment 1, millet vs. no millet in experiment 2), alternating choices (same arm as before vs. different) and pattern (only experiment 2, dots vs. stripes). The analysis of alternating was done by labelling the choices according to whether the arm chosen in this trial was also the arm chosen in the trial before.The first day of both weeks, respectively, were excluded from this labelling.

The results from all mice were then used for significance testing: To test for normal distribution, the Shapiro–Wilk test was performed in R. The data was normal distributed (p > 0.05); therefore, a t-test was used to compare the percentages of the mice with a random chance level of 0.5. In all statistical tests, significance level was set to 0.05, and result values are given as mean and standard deviation. (For more details on the analysis, especially with regard to the passive habituation of experiment 1, see Supplements.)

## 3 Results Experiment 1

### 3.1 Passive Habituation

After one week, all mice except for one visited the T-maze frequently. After nine days, all thirteen mice did so. As in retrospect was noted that the RFID registration system might have worked inaccurate (although this was not the case when tested before), some passages might have been missed. Thus, habituation might have been even better than noted.

### 3.2 Trial Duration and Inter Trial Interval in the T-Maze

In most cases, mice self-initiated the trials: Only in two out of 143 trials (habituation trials and miss-recorded trials included), a mouse did not start the trial by itself within the set start time and had to be guided by tube handling into the maze.

Habituation trials included a visit in both arms. From the time point when the mice entered the T-maze to the time point when the mice had crossed the virtual line in both arms, on average 17.2 ± 11.6 s passed (minimum: 7.5 s, maximum: 45.5 s). For the preference test trials, average duration was 4.46 ± 2.93 s (minimum 1.25 s, maximum: 25.9 s). Note that in this experimental setup, the way back to the start cage was not blocked so mice could return to the start cage and later on re-visit the maze. The numbers given here are only from those times when a mouse entered the maze and actually crossed one of the virtual lines. Mean intertrial interval (ITI), including cleaning time of the maze and the time until the mouse decided to enter the maze once again, was 204.9 ± 81.8 s (= 3.4 min), ranging from a minimum of 137 s to a maximum of 506 s.

### 3.3 Preference Testing

It was not possible to compare the intake of the offered fluid droplet between apple juice and almond milk on the basis of the video recordings as it was only detectable for the opaque almond milk whether it disappeared. Still, we assessed whether the animals spent some time investigating the droplet (licking or intensely sniffing it). This was observed in 74 of 139 trials (including only one time during a habituation trial), representing barely more than half of the trials. 75.67 % of these observed behaviours were performed towards an almond milk droplet.

Comparing the choices of the mice for the arm with apple juice or the arm with almond milk, mice chose in 52.8 ± 9.9 % of the trials the arm with almond milk. This indicates no preference (t = 1.028, df = 12, p = 0.3242, see Fig. 3). Mice showed also no side preference: The left arm was chosen on average in 49.5 ± 14.1 % of trials (t = -0.13145, df = 12, p = 0.8976). As the T-maze test is often used to test for spontaneous alternation (Deacon2006), we then analysed the data with regard to alternating choices. Indeed, mice chose in 64.4 ± 13.5 % of trials the arm which they did not choose during the last trial (t = 3.8442, df = 12, p < 0.003).

**Fig. 3:**
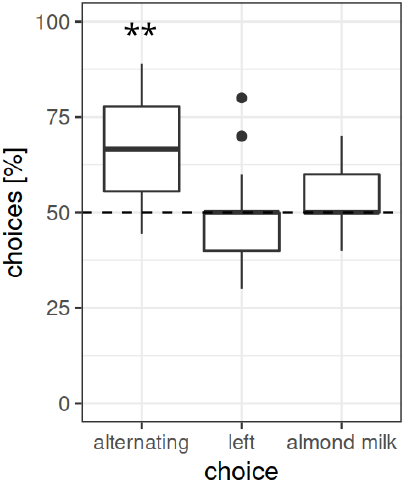
Percentage of choices for the arm not visited in the preceding trial (alternating), the left arm and the arm containing almond milk. Thirteen female mice chose ten times (5 per day) between an arm containing the odour and a 20 *µ*l droplet of almond milk or apple juice. Presentation side was randomized across the group, and switched after trial seven. ** p < 0.01

## 4 Results Experiment 2

### 4.1 Active Habituation

Mice were familiar with automated doors from previous experiments. However, in this new setup they seemed to experience the door as something new, so that on day one of habituation, only one mouse went into the maze on its own. Nevertheless, on the fifth day of habituation all mice went into the maze by themselves within the time frame of three minutes.

### 4.2 Trial Duration and Inter Trial Interval in the T-Maze

Time spent by the mice in the start cage before entering the maze ranged between 1.7 and 159.5 s (on average 21.27 ± 22.71 s). Inside the maze, the mice took only 3.6 ± 1.7 s to make a choice and enter one of the goal arms far enough to cross the virtual line (min: 1.4 s, max: 14.5 s). There, mice spent about 47.4 ± 33.09 s in the arm before entering the provided tube. After preparing the arms again for the next trial, the mouse was returned to the start cage. This intertrial interval lasted on average 19.4 ± 8.8 s (min: 4 s, max: 107 s, caused by an error during the preparation), measuring the time between the mice being taken out of the arm and starting the new trial. Including the time between making the choice and leaving the arm would add the approximately 47 s spent in the goal arm.

### 4.3 Preference Testing

In week 1 (two trials per day, side change after trial six), mice chose in 43.3 ± 8.9 % the arm containing millet, which meant that they significantly preferred the arm without it (t = -2.6018, df = 11, p < 0.05, see Fig. 4a). There was no side preference (left arm chosen in 50.0 ± 12.8 %, t = 0, df = 11, p = 1.00) and no pattern preference (dots chosen in 55.0 ± 10 %, t = 1.7321, df = 11, p = 0.11). However, mice also significantly alternated between arms (63.9 ± 15.8 %, t = 3.0446, df = 11, p < 0.05).

**Fig. 4:**
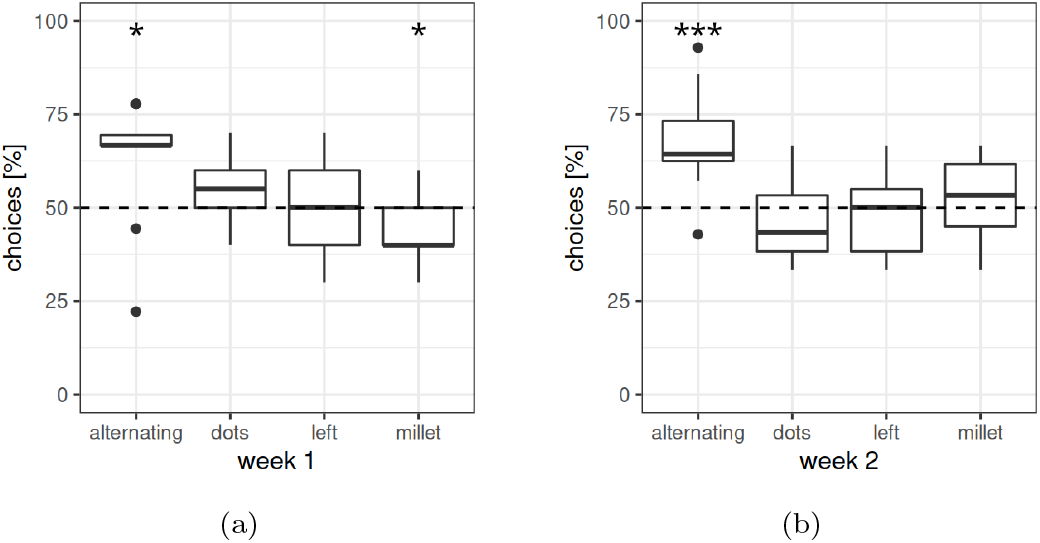
Percentage of choices for the arm not visited in the preceding trial (alternating), the arm marked with dots, the left arm and the arm containing almond milk. One group of twelve female mice chose between an arm containing bedding mixed with millet and an arm only containing bedding. Presentation side and pattern (dots or stripes) was randomized across the group. (a) In week 1, two trials were performed per day (ten in total), and after trial six, presentation side was switched. (b) In week 2, three trials were performed per day (fifteen in total), and presentation side was kept as last used in week 1. * p < 0.05, *** p < 0.001

In week 2 (three trials per day, no side change), mice chose in 53.4 ±-11.4 % the arm containing millet (t = 1.0155, df = 11, p = 0.33, see Fig. 4b). There was no side preference (left: t = -0.6603, df = 11, 47.8 ± 11.7 %, p = 0.52) or pattern preference (dots: 45.6 ± 10.9 %, t = -1.4062, df = 11, p = 0.19) but mice significantly alternated between trials (67.9 ± 13.4 %, t = 4.5993, df = 11, p < 0.001). When looking at the individual trials (see Fig. 5), percentage of alternation was especially apparent in the second and third trial but not in the first, which was compared to the last trial on the day before (week 1: trial 1 52.1 *±* 19.8 %, trial 2: 73.3 *±* 27.4 %; week 2: trial 1 43.8 *±* 24.1 %, trial 2 81.7 ± 19.9 %, trial 3 73.3 *±* 17.8 %).

**Fig. 5:**
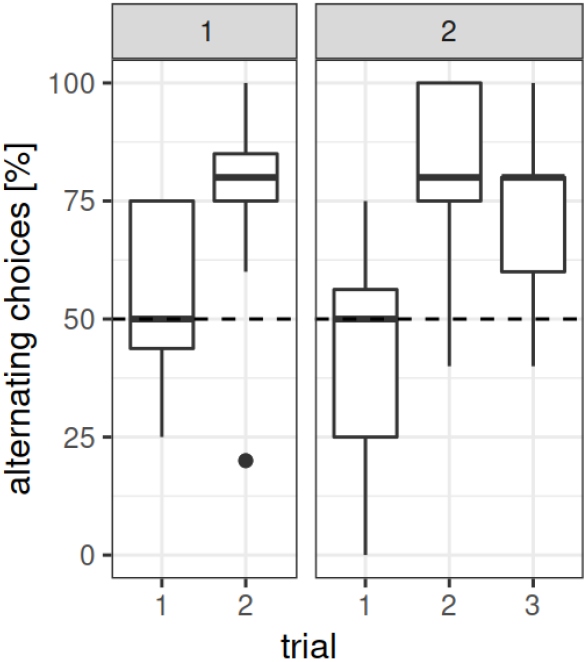
Percentage of choices for the arm not visited in the preceding trial (alternating) across trials for week 1 (left, two trials per day) and week 2 (right, three trials per day). One group of twelve female mice chose between an arm containing bedding mixed with millet and an arm only containing bedding. Presentation side and pattern (dots or stripes) was randomized across the group. In week 1, two trials were performed per day (ten in total), and after trial six, presentation side was switched. In week 2, three trials were performed per day (fifteen in total), and presentation side was kept as last used in week 1.

## 5 Discussion

### 5.1 Habituation

Mice took on average about 10 s (experiment 1) or 4 s (experiment 2) to make a choice after starting the trial. This implies that mice were well habituated: As Deacon and Rawlins (2006) describe, a trial duration longer than two minutes can indicate insufficient habituation, and here, mice were much faster. However, the two minutes Deacon and Rawlins (2006) use as a bench mark, usually include the time from placing the animal in the start area of the maze to the actual choice (Deacon and Rawlins 2006). We here provided the animal the opportunity to self-initiate the test, which probably conduced to a shorter trial time because trials started apparently when the animal itself was motivated.

However, it is possible that animals were not habituated enough for the preference test itself: Judging on the basis of their behaviour in experiment 1, mice tested the fluid drop only in half of the trials. This might be an indication for insufficient habituation, as during pre-tests before the second experiment, mice fed on millet in an unfamiliar surrounding only after several sessions of habituating to it. In addition, we observed during the pre-tests that millet was consumed more willingly in general than almond milk. Therefore, in experiment 2, one week of active instead of passive habituation to the T-maze was conducted, and we used millet as a reward. Here, all mice fed on the millet when choosing the respective arm. Thus, feeding behaviour in the maze seems to be influenced by both the habituation method and the type of reward.

### 5.2 Lack of preference or reward-aimed behaviour

In preparation of experiment 1, when offering the two fluids in the home cage for habituation, the twelve mice as a group drank nearly 500 ml in 24 h of the almond milk in 24 h, whereas they drank only about 200 ml of the provided apple juice. This implies a strong preference. However, no fluid preference was found in the T-maze preference test, and there are various possible reasons for it.

One explanation might be a missing cue on where to find the almond milk. In the first experiment, in addition to spatial information (at least during the first seven trials) an odour cue was provided. However, between the trials, the maze was cleaned with ethanol to erase odour cues. This was done because intramaze odour cues of previous decisions might influence the next choice (rats: Means et al. 1992). Nevertheless, the ethanol itself might have left an odour, masking the olfactory cue of almond milk and apple juice. We investigated this theory by not cleaning the maze in-between mice in experiment 2. Although we did not provide an additional olfactory cue on a cellulose sheet as in experiment 1, it can be assumed that the options (millet or no millet) naturally include an olfactory cue. In addition, a visual cue (wall pattern) and a spatial cue (no side change in week 2) were provided. Thus, mice should have had the possibility to learn which of the two arms was the rewarded one. However, this also did not lead to a preference for the rewarded arm. It is not likely that this occurred because mice were sated on millet after consuming three times 0.1 g because in another experiment in our research group, mice received about 0.8 g millet per day and were still willing to work for it.

As the setup of experiment 2 is in general similar to simple learning tests (operant conditioning, learning the relationship between behaviour and its outcome), mice should be able to learn the position of the millet. For optimal foraging, animals should adopt in this scenario the win – stay/lose – shift strategy, meaning that they should stay (or return to) where they found food before and change position when they did not find food (Shettleworth 2010). Thus, we would have expected the mice to perform the above mentioned strategy. However, not all animals follow one of those strategies (e.g., chickadees are known to neither use win – stay nor win – shift strategies(Guitar et al. 2017), and noisy miner birds use both strategies, depending on the reward type (Sulikowski and Burke 2010)). Also, when a relationship between behaviour and a reward is learned, it can take time to reverse it (Mackintosh et al. 1968). Nevertheless, here we used mice naive to the T-maze, so they should not have formed a previous association which could interfere. It seems we observed a similar result here as described in the study by Locurto et al. (2002), in which offspring of a C57BL/6 and DBA/2J cross easily learned the win – shift strategy but did not exceed chance levels when requested to perform win – stay (Locurto et al. 2002; also Locurto 2005). This is in contrast to other studies which successfully report using the T-maze for discrimination tests (spatial or visual) and the learning of the win – stay strategy (Lione et al. 1999; Granholm et al. 2000; Belzung et al. 2001).

One premise for showing the win – stay strategy would be remembering what was done before to find food. As trials were performed on multiple days, remembering the last choice made on the day before might not have been possible for the mice and the first choice was always based on chance. Thus, with only two trials per day, a preference might have therefore been disguised in week 1 of experiment 2. However, in week 2, there were always three trials per day. This means even if the mice had not remembered the position of the millet from the day before, after two trials of sampling, the third trial should have been based on a preference. As a result, it could have been expected that a) all third trials were made towards the millet arm, and b) the preference for millet in total was at least in 2/3 of the trials. However, this was not the case as alternation levels in the third trial were similar to the second, and portion of chosen millet arms was about 1/2.

Another factor that might prevent manifestation of the win – stay strategy could be that mice found the reward already lying in the arm instead of receiving a reward when entering the arm (experiencing the arm as empty but then getting food). As a result, when leaving the arm after eating all the millet, they might have memorised this arm as empty of millet. This might correspond to the findings of Herrmann et al. (1982), who performed a three-table task with rats: After some exploration time in the apparatus, rats received their reward on one of the three tables. If they were allowed to completely feed on the food, rats were able to learn win – shift but not win – stay. If they were only allowed to feed partially, rats win – stay behaviour was faster shown than win – shift (Herrmann et al. 1982). This indicates that the animals remember whether the feeding place was emptied or not and could explain why mice did seldom return to the arm in which they experienced food beforehand. Thus, one way of improving the procedure could be to allow only partial feeding in the goal arms. Another possibility would be to “show” the mice that the feeding place is refilled. This would be inspired by conditioned place preference and the study of Goltseker and Barak (2018): Here, conditioned place aversion was only induced when mice where placed in an empty compartment first, and then experienced the onset of the aversive stimulus (in this case: cold water flooding). Conditioned place aversion was not induced when the mice were placed in an already flooded compartment (Goltseker and Barak 2018). This implies that timing plays an important role for association formation. However, in the case of the T-maze as a preference test, providing the good after the mouse enters the arm would request more experimenter interaction (and therefore, more possible experimenter influence and bias) or a thorough automation. It is also important to keep in mind that in this case the odour stimulus would be erased.

Another factor that might prevent manifestation of the win – stay strategy might be that mice had other motivations than to search for a preferred fluid or a food reward in the maze. As described by Dixon et al. (2013), additional motivations can influence behaviour and the results of preference tests. Here, results of the conditioned place preference test were undermined by the motivation of the birds to search for food or to stay in the more familiar compartment (the one experienced last) (Dixon et al. 2013). However, it cannot be said that mice showed no preference in their behaviour at all. Instead, they showed a clear preference for the arm which they had not visited during the last trial, a behaviour which is called “spontaneous alternation”.

### 5.3 Influences on spontaneous alternation

Spontaneous alternation behaviour is a common phenomenon in the T-maze (Deacon and Rawlins 2006; Sharma et al. 2010b; Dember and Fowler 1958). There are many theories on the factors that influence spontaneous alternation (also reviewed in Richman et al. 1986) and in the following we will shortly discuss some of them.

Apparently, orientation of the maze arms influences spontaneous alternation (Dember and Fowler 1958). I.e., alternation decreases when both arms lead towards the same goal. Also, if the arms are positioned not opposite to each other but parallel, spontaneous alternation is reduced (Novak et al. 2016b; Novak et al. 2016a).

For rats it was shown that external stimuli (spatial cues) seem to be more important for alternation than proprioceptive cues, i.e., the body’s turning response to the left or right (non-spatial cues) (Montgomery 1952). A maze rich of spatial extramaze cues reinforces alternation (Lennartz 2008) but spatial intramaze cues (e.g., patterns on the arm walls) have stronger influence on alternation than extramaze cues (Walker et al. 1955). In mice, however, influence of spatial and non-spatial cues seems to differ between strains. C57BL/6J, for example, did not exceed chance level in a spatial task using extramaze cues but were slightly better in a non-spatial proprioceptive task (left vs. right turn). BALB/cByJ, on the other hand, performed well in both tasks (Crusio et al. 1990).

In our experiments, we used C57BL/6J mice, and we provided several cues: In both experiment for most of the trials (except for those after the side change) the spatial as well as the non-spatial cue were the same. In addition, we provided an olfactory (experiment 1) and a visual (experiment 2) intramaze cue, without notable changes in alternation behaviour. Moreover, in experiment 2 odour trails from previous trials could have functioned as a cue, in which the maze was not disinfected between the trials. It is evident that the mice did not use any of the provided cues to choose the supposedly more rewarding arm. However, the cues might have influenced alternation as it is discussed that animals might be driven to explore the stimulus which is less familiar, i.e., to which they were not exposed last (Richman et al. 1986). For example, mutant mice showing no or reduced habituation also showed no spontaneous alternation behaviour in the T-maze (Lalonde et al. 1986). This implies that there are indeed additional motivations during the test which interfere with the motivation to gain food reward.

In general, spontaneous alternation behaviour seems also to be intertrial interval (ITI) time (and thus, memory) dependent. However, regarding which ITIs support spontaneous alternation and which not, the literature is mixed. Durantou et al. (1989) report high alternation rates for ITIs of 30 s in rats (Durantou et al. 1989), whereas in the T-maze protocol for appetitive position discrimination for both rats and mice by Deacon 2006, it is stated that in general ITIs shorter than 10 min lead to spontaneous alternation. In the review of Dember and Fowler 1958 focusing on alternation in rats, it is described that spontaneous alternation was observed for ITIs shorter than 2 min but in some studies also up to 60 min. Here, in experiment 2, ITI was about 19 s but never longer than 2 min, and in experiment 1, ITI lasted about 3.5 min. This fits to the description made by Deacon (2006) for mice. We can also confirm that for long ITIs alternation drops to chance level (Deacon 2006; Durantou et al. 1989): Comparing alternation proportions of individual trials for experiment 2 revealed less alternation behaviour during the first trial of each day. Thus, the last choice of the day before (with an ITI > 21 h) seems not to be relevant for the first choice. However, one of the problems of comparing the influence of it is might be that studies use different definitions what they exactly consider to be the intertrial interval. For example, Locurto (2005) regards the ITI as the time between two trials but with one trial consisting of two forced choice trails and one free choice trial, meaning the time between the forced choice and the free choice trials is not considered (Locurto 2005). It is also discussed whether food reward itself influences alternation behaviour, and if so, under which circumstances. Apparently, at least in rats alternation levels are reduced with increasing food deprivation (Richman et al. 1986). Returning to the topic of the foraging strategies, this implies that rats switch to win – stay strategy only when the motivation to gain food is high enough. In other words: Below a specific food deprivation level, the motivation to explore what was not experienced in the preceding trial might be higher than the motivation to gain food (Richman et al. 1986). This exploration behaviour could be driven by additional needs, for example, search for shelter (Pilz et al. 2020) or an escape out of the maze.

In this context, it has also to be kept in mind that it was shown already in the 1960s that conditioned stimuli are not equally effective for all kinds of unconditioned stimuli, for example, gustatory and olfactory stimuli are more easily associated with internal discomfort than audio-visual stimuli (Garcia and Koelling 1966). This learning phenomenon is probably caused by an evolutionary advantage of facilitated association of specific stimuli. In a similar manner, evolution might have favoured learning mechanisms which cause mice to prefer the win – shift strategy under *ad libitum* food conditions and the win – stay strategy under food restricted conditions. Thus, asking the mice to choose a food rewarded arm over an empty arm might be a completely different question under different feeding conditions.

It has to be mentioned that an additional important factor for alternation seems to be fear or stress. Under the key word “optimal arousal theory” multiple studies can be found, which investigate the effect of a mild stressor (open field test), food shock or water presence (water-escape T-maze instead of dry T-maze) on the alternating behaviour (rats: Means 1988; Comer and Means 1989; mice: Mitchell et al. 1984; Mitchell et al. 1985; Bats et al. 2001). In general, this theory suggests that individuals seek the optimal arousal, which is shaped in an upside-down u-curve. Thus, when an animal is not aroused it would seek something arousing, for example, a less familiar environment. When the animal is already “too much” aroused (behind the peak of the curve), however, it would seek the less arousing stimuli, meaning a more familiar environment. This theory tries to explain why after experiencing a mild stressor, mice perseverated their choices instead of alternating (Bats et al. 2001). In an other experiment of Mitchell et al. (1984), mice received food shocks while running through the maze, which terminated when they chose the goal arm which they did not choose in the preceding trial (alternated) but continued when they chose the same goal arm again (perseverated). However, mice did not learn this association and kept on perseverating. Mitchell et al. (1984) called it the “punishment paradox” (Mitchell et al. 1984). (It has to be noted that in this experiment, the shock continued in the wrong arm. Mice would probably behave differently if the shock was administered only when entering the wrong arm.) Transferring these observations to our experiments, we could conclude that the procedure before and during the T-maze was probably not stressful as our mice did not perseverate but alternate. What we observed was rather the “alternating paradox”, meaning alternating although perseverating was reinforced.

## 6 Conclusion

It is obvious that T-maze as used in this setup was not suitable to investigate preference or reward-aimed learning in C57BL/6J mice. Instead, mice alternated their choices in 60-70 % of the trials. Although the main reason behind this alternation behaviour remains unclear, we can at least validate the statement by Deacon and Rawlins 2006 that well habituated animals run the T-maze alternation test also without food restriction. It might be possible to increase performance by imposing deprivation on the animals. However, as we were interested in preference under un-restrained conditions, we deem the T-maze not suitable for our research question.

## Ethical Approval

All experiments were approved by the Berlin state authority, Landesamt für Gesundheit und Soziales, under license No. G 0182/17 and were in accordance with the German Animal Protection Law (TierSchG, TierSchVersV).

The second experiment was preregistered at the Animal Study Registry (DOI: 10.17590/asr.0000213).

## Authors Contribution

Conceptualization: Anne Habedank Pia Kahnau Lars Lewejohann

Methodology: Anne Habedank

Formal analysis and investigation: Anne Habedank

Writing - original draft preparation: Anne Habedank

Writing - review and editing: Anne Habedank Lars Lewejohann Pia Kahnau

Funding acquisition: Lars Lewejohann

Supervision: Lars Lewejohann

## Acknowledgements

The authors thank the animal caretakers, especially Carola Schwarck, for their support in the animal husbandry.

## References

Acosta J, Bussi IL, Esquivel M, Höcht C, Golombek DA, Agostino PV (2020) Circadian modulation of motivation in mice. Behavioural Brain Research 382:112471, DOI 10.1016/j.bbr.2020.112471, URL https://doi.org/10.1016/j.bbr.2020.112471

Bats S, Thoumas J, Lordi B, Tonon M, Lalonde R, Caston J (2001) The effects of a mild stressor on spontaneous alternation in mice. Behavioural Brain Research 118(1):11–15, DOI 10.1016/s0166-4328(00)00285-0, URL https://doi.org/10.1016/s0166-4328(00)00285-0

Belzung C, Chapillon P, Lalonde R (2001) The effects of the lurcher mutation on object localization, t-maze discrimination, and radial arm maze tasks. Behavior Genetics 31(2):151–155, DOI 10.1023/a:1010269126295, URL https://doi.org/10.1023/a:1010269126295

Buckley LA, Sandilands V, Tolkamp BJ, D’Eath RB (2011) Quantifying hungry broiler breeder dietary preferences using a closed economy t-maze task. Applied Animal Behaviour Science 133(3-4):216–227, DOI 10.1016/j.applanim.2011.06.003, URL https://doi.org/10.1016/j.applanim.2011.06.003

Comer TR, Means LW (1989) Overcoming unlearned response biases: delayed escape following errors facilitates acquisition of win-stay and win-shift working memory water-escape tasks in rats. Behavioral and Neural Biology 52(2):239–250, DOI 10.1016/s0163-1047(89)90355-5, URL https://doi.org/10.1016/s0163-1047(89)90355-5

Correa M, Pardo M, Bayarri P, López-Cruz L, Miguel NS, Valverde O, Ledent C, Salamone JD (2015) Choosing voluntary exercise over sucrose consumption depends upon dopamine transmission: effects of haloperidol in wild type and adenosine a2ako mice. Psychopharmacology 233(3):393–404, DOI 10.1007/s00213-015-4127-3, URL https://doi.org/10.1007/s00213-015-4127-3

Crusio W, Bertholet JY, Schwegler H (1990) No correlations between spatial and non-spatial reference memory in a t-maze task and hippocampal mossy fibre distribution in the mouse. Behavioural Brain Research 41(3):251–259, DOI 10.1016/0166-4328(90)90112-r, URL https://doi.org/10.1016/0166-4328(90)90112-r

Cunningham CL, Patel P, Milner L (2006) Spatial location is critical for conditioning place preference with visual but not tactile stimuli. Behavioral Neuroscience 120(5):1115–1132, DOI 10.1037/0735-7044.120.5.1115, URL https://doi.org/10.1037/0735-7044.120.5.1115

Cunningham PJ, Kuhn R, Reilly MP (2015) A within-subject between-apparatus comparison of impulsive choice: T-maze and two-lever chamber. Journal of the Experimental Analysis of Behavior 104(1):20–29, DOI 10.1002/jeab.159, URL https://doi.org/10.1002/jeab.159

Cutuli D, Caporali P, Gelfo F, Angelucci F, Laricchiuta D, Foti F, Bartolo PD, Bisicchia E, Molinari M, Vecchioli SF, Petrosini L (2015) Pre-reproductive maternal enrichment influences rat maternal care and offspring developmental trajectories: behavioral performances and neuroplasticity correlates. Frontiers in Behavioral Neuroscience 9, DOI 10.3389/fnbeh.2015.00066, URL https://doi.org/10.3389/fnbeh.2015.00066

Dawkins M (1977) Do hens suffer in battery cages? environmental preferences and welfare. Animal Behaviour 25:1034–1046, DOI 10.1016/0003-3472(77)90054-9, URL https://doi.org/10.1016/0003-3472(77)90054-9

Deacon RMJ (2006) Appetitive position discrimination in the t-maze. Nature Protocols 1(1):13–15, DOI 10.1038/nprot.2006.3, URL https://doi.org/10.1038/nprot.2006.3

Deacon RMJ, Rawlins JNP (2006) T-maze alternation in the rodent. Nature Protocols 1(1):7–12, DOI 10.1038/nprot.2006.2, URL https://doi.org/10.1038/nprot.2006.2

Dember WN, Fowler H (1958) Spontaneous alternation behavior. Psychological Bulletin 55(6):412–428, DOI 10.1037/h0045446, URL https://doi.org/10.1037/h0045446

Dember WN, Kleinman R (1973) Cues for spontaneous alternation by gerbils. Animal Learning & Behavior 1(4):287–289, DOI 10.3758/bf03199253, URL https://doi.org/10.3758/bf03199253

Denk F, Walton ME, Jennings KA, Sharp T, Rushworth MFS, Bannerman DM (2004) Differential involvement of serotonin and dopamine systems in cost-benefit decisions about delay or effort. Psychopharmacology 179(3):587–596, DOI 10.1007/s00213-004-2059-4, URL https://doi.org/10.1007/s00213-004-2059-4

Dixon LM, Sandilands V, Bateson M, Brocklehurst S, Tolkamp BJ, D’Eath RB (2013) Conditioned place preference or aversion as animal welfare assessment tools: Limitations in their application. Applied Animal Behaviour Science 148(1-2):164–176, DOI 10.1016/j.applanim.2013.07.012, URL https://doi.org/10.1016/j.applanim.2013.07.012

Dudchenko PA (2004) An overview of the tasks used to test working memory in rodents. Neuroscience & Biobehavioral Reviews 28(7):699–709, DOI 10.1016/j.neubiorev.2004.09.002, URL https://doi.org/10.1016/j.neubiorev.2004.09.002

Durantou F, Cazala P, Jaffard R (1989) Intertrial interval dependent effect of lateral hypothalamic stimulation on spontaneous alternation behavior in a t-maze. Physiology & Behavior 46(2):253–258, DOI 10.1016/0031-9384(89)90264-3, URL https://doi.org/10.1016/0031-9384(89)90264-3

Fitchett AE, Barnard CJ, Cassaday HJ (2006) There’s no place like home: Cage odours and place preference in subordinate CD-1 male mice. Physiology & Behavior 87(5):955–962, DOI 10.1016/j.physbeh.2006.02.010, URL https://doi.org/10.1016/j.physbeh.2006.02.010

Friard O, Gamba M (2016) BORIS: a free, versatile open-source event-logging software for video/audio coding and live observations. Methods in Ecology and Evolution 7(11):1325–1330, DOI 10.1111/2041-210x.12584, URL https://doi.org/10.1111/2041-210x.12584

Fujita M, Tanimura T (2011) Drosophila evaluates and learns the nutritional value of sugars. Current Biology 21(9):751–755, DOI 10.1016/j.cub.2011.03.058, URL https://doi.org/10.1016/j.cub.2011.03.058

Garcia J, Koelling RA (1966) Relation of cue to consequence in avoidance learning. Psychonomic Science 4(1):123–124, DOI 10.3758/bf03342209, URL https://doi.org/10.3758/bf03342209

Gerlai R (1998) A new continuous alternation task in t-maze detects hippocampal dysfunction in mice. Behavioural Brain Research 95(1):91–101, DOI 10.1016/s0166-4328(97)00214-3, URL https://doi.org/10.1016/s0166-4328(97)00214-3

Goltseker K, Barak S (2018) Flood-conditioned place aversion as a novel non-pharmacological aversive learning procedure in mice. Scientific Reports 8(1), DOI 10.1038/s41598-018-25568-5, URL https://doi.org/10.1038/s41598-018-25568-5

Gouveia K, Hurst JL (2017) Optimising reliability of mouse performance in behavioural testing: the major role of non-aversive handling. Scientific Reports 7(1), DOI 10.1038/srep44999, URL https://doi.org/10.1038/srep44999

Granholm ACE, Sanders LA, Crnic LS (2000) Loss of cholinergic phenotype in basal forebrain coincides with cognitive decline in a mouse model of down’s syndrome. Experimental Neurology 161(2):647–663, DOI 10.1006/exnr.1999.7289, URL https://doi.org/10.1006/exnr.1999.7289

Guariglia SR, Chadman KK (2013) Water t-maze: A useful assay for determination of repetitive behaviors in mice. Journal of Neuroscience Methods 220(1):24–29, DOI 10.1016/j.jneumeth.2013.08.019, URL https://doi.org/10.1016/j.jneumeth.2013.08.019

Guitar NA, Strang CG, Course CJ, Sherry DF (2017) Chickadees neither win-shift nor win-stay when foraging. Animal Behaviour 133:73–82, DOI 10.1016/j.anbehav.2017.09.011, URL https://doi.org/10.1016/j.anbehav.2017.09.011

Hernandez-Lallement J, van Wingerden M, Marx C, Srejic M, Kalen-scher T (2015) Rats prefer mutual rewards in a prosocial choice task. Frontiers in Neuroscience 8, DOI 10.3389/fnins.2014.00443, URL https://doi.org/10.3389/fnins.2014.00443

Herrmann T, Bahr E, Bremner B, Ellen P (1982) Problem solving in the rat: Stay vs. shift solutions on the three-table task. Animal Learning & Behavior 10(1):39–45, DOI 10.3758/bf03212044, URL https://doi.org/10.3758/bf03212044

Hieu BTN, Anh NTN, Audira G, Juniardi S, Liman RAD, Villaflores OB, Lai YH, Chen JR, Liang ST, Huang JC, Hsiao CD (2020) Development of a modified three-day t-maze protocol for evaluating learning and memory capacity of adult zebrafish. International Journal of Molecular Sciences 21(4):1464, DOI 10.3390/ijms21041464, URL https://doi.org/10.3390/ijms21041464

Hurst JL, West RS (2010) Taming anxiety in laboratory mice. Nature Methods 7(10):825–826, DOI 10.1038/nmeth.1500, URL https://doi.org/10.1038/nmeth.1500

Kirkden RD, Pajor EA (2006) Using preference, motivation and aversion tests to ask scientific questions about animals’ feelings. Applied Animal Behaviour Science 100(1-2):29–47, DOI 10.1016/j.applanim.2006.04.009, URL https://doi.org/10.1016/j.applanim.2006.04.009

Koch CE, Begemann K, Kiehn JT, Griewahn L, Mauer J, Hess ME, Moser A, Schmid SM, Brüning JC, Oster H (2020) Circadian regulation of hedonic appetite in mice by clocks in dopaminergic neurons of the VTA. Nature Communications 11(1), DOI 10.1038/s41467-020-16882-6, URL https://doi.org/10.1038/s41467-020-16882-6

Lalonde R, Botez MI, Boivin D (1986) Spontaneous alternation and habituation in a t-maze in nervous mutant mice. Behavioral Neuroscience 100(3):350–352, DOI 10.1037/0735-7044.100.3.350, URL https://doi.org/10.1037/0735-7044.100.3.350

Leenaars CH, van der Mierden S, Durst M, Goerlich-Jansson VC, Ripoli FL, Keubler LM, Talbot SR, Boyle E, Habedank A, Jirkof P, Lewejohann L, Gass P, Tolba R, Bleich A (2019) Measurement of corticosterone in mice: a protocol for a mapping review. Laboratory Animals 54:26–32, DOI 10.1177/0023677219868499, URL https://doi.org/10.1177/0023677219868499

Lennartz RC (2008) The role of extramaze cues in spontaneous alternation in a plus-maze. Learning & Behavior 36(2):138–144, DOI 10.3758/lb.36.2.138, URL https://doi.org/10.3758/lb.36.2.138

Lione LA, Carter RJ, Hunt MJ, Bates GP, Morton AJ, Dunnett SB (1999) Selective discrimination learning impairments in mice expressing the human huntington’s disease mutation. The Journal of Neuroscience 19(23):10428–10437, DOI 10.1523/jneurosci.19-23-10428.1999, URL https://doi.org/10.1523/jneurosci.19-23-10428.1999

Locurto C (2005) Further evidence that mice learn a win-shift but not a win-stay contingency under water-escape motivation. Journal of Comparative Psychology 119(4):387–393, DOI 10.1037/0735-7036.119.4.387, URL https://doi.org/10.1037/0735-7036.119.4.387

Locurto C, Emidy C, Hannan S (2002) Mice (mus musculus) learn a win-shift but not a win-stay contingency under water escape motivation. Journal of Comparative Psychology 116(3):308–312, DOI 10.1037/0735-7036.116.3.308, URL https://doi.org/10.1037/0735-7036.116.3.308

Lohninger S, Strasser A, Bubna-Littitz H (2001) The effect of l-carnitine on t-maze learning ability in aged rats. Archives of Gerontology and Geriatrics 32(3):245–253, DOI 10.1016/s0167-4943(01)00097-8, URL https://doi.org/10.1016/s0167-4943(01)00097-8

Mackintosh NJ, Mcgonigle B, Holgate V (1968) Factors underlying improvement in serial reversal learning. Canadian Journal of Psychology/Revue canadienne de psychologie 22(2):85–95, DOI 10.1037/h0082753, URL https://doi.org/10.1037/h0082753

Mayeux-Portas V, File SE, Stewart CL, Morris RJ (2000) Mice lacking the cell adhesion molecule thy-1 fail to use socially transmitted cues to direct their choice of food. Current Biology 10(2):68–75, DOI 10.1016/s0960-9822(99)00278-x, URL https://doi.org/10.1016/s0960-9822(99)00278-x

Means LW (1988) Rats acquire win-stay more readily than win-shift in a water escape situation. Animal Learning & Behavior 16(3):303–311, DOI 10.3758/bf03209081, URL https://doi.org/10.3758/bf03209081

Means LW, Alexander SR, O’Neal MF (1992) Those cheating rats: male and female rats use odor trails in a water-escape “working memory” task. Behavioral and Neural Biology 58(2):144–151, DOI 10.1016/0163-1047(92)90387-j, URL https://doi.org/10.1016/0163-1047(92)90387-j

Mitchell D, Koleszar A, Scopatz RA (1984) Arousal and t-maze choice behavior in mice: A convergent paradigm for neophobia constructs and optimal arousal theory. Learning and Motivation 15(3):287–301, DOI 10.1016/0023-9690(84)90024-9, URL https://doi.org/10.1016/0023-9690(84)90024-9

Mitchell D, Osborne EW, O’Boyle MW (1985) Habituation under stress: Shocked mice show nonassociative learning in a t-maze. Behavioral and Neural Biology 43(2):212–217, DOI 10.1016/s0163-1047(85)91387-1, URL https://doi.org/10.1016/s0163-1047(85)91387-1

Montgomery KC (1952) A test of two explanations of spontaneous alternation. Journal of Comparative and Physiological Psychology 45(3):287–293, DOI 10.1037/h0058118, URL https://doi.org/10.1037/h0058118

Moy SS, Nadler JJ, Young NB, Nonneman RJ, Segall SK, Andrade GM, Crawley JN, Magnuson TR (2008) Social approach and repetitive behavior in eleven inbred mouse strains. Behavioural Brain Research 191(1):118–129, DOI 10.1016/j.bbr.2008.03.015, URL https://doi.org/10.1016/j.bbr.2008.03.015

Nasrawi CW, Pangborn RM (1990) Temporal effectiveness of mouth-rinsing on capsaicin mouth-burn. Physiology & Behavior 47(4):617–623, DOI 10.1016/0031-9384(90)90067-e, URL https://doi.org/10.1016/0031-9384(90)90067-e

Novak J, Bailoo JD, Melotti L, Würbel H (2016a) Effect of cage-induced stereotypies on measures of affective state and recurrent perseveration in CD-1 and c57bl/6 mice. PLOS ONE 11(5):e0153203, DOI 10.1371/journal.pone.0153203, URL https://doi.org/10.1371/journal.pone.0153203

Novak J, Stojanovski K, Melotti L, Reichlin TS, Palme R, Würbel H (2016b) Effects of stereotypic behaviour and chronic mild stress on judgement bias in laboratory mice. Applied Animal Behaviour Science 174:162–172, DOI 10.1016/j.applanim.2015.10.004, URL https://doi.org/10.1016/j.applanim.2015.10.004

Nunes AC, da Luz Mathias M, Ganem G (2009) Odor preference in house mice: influences of habitat heterogeneity and chromosomal incompatibility. Behavioral Ecology 20(6):1252–1261, DOI 10.1093/beheco/arp122, URL https://doi.org/10.1093/beheco/arp122

Patterson-Kane EG, Harper DN, Hunt M (2001) The cage preferences of laboratory rats. Laboratory Animals 35(1):74–79, DOI 10.1258/0023677011911390, URL https://doi.org/10.1258/0023677011911390

Pennycuik P, Cowan R (1990) Odor and food preferences of house mice, mus-musculus. Australian Journal of Zoology 38(3):241, DOI 10.1071/zo9900241, URL https://doi.org/10.1071/zo9900241

Pilz P, Mück F, Walter M (2020) Learning behaviour of mice in multiple t-versus y-mazes. Abstract Booklet of the 15th Annual Meeting of the Ethological Society

Pioli EY, Gaskill BN, Gilmour G, Tricklebank MD, Dix SL, Bannerman D, Garner JP (2014) An automated maze task for assessing hippocampus-sensitive memory in mice. Behavioural Brain Research 261:249–257, DOI 10.1016/j.bbr.2013.12.009, URL https://doi.org/10.1016/j.bbr.2013.12.009

van der Plasse G, Fors SSBML, Meerkerk DTJ, Joosten RNJMA, Uylings HBM, Feenstra MGP (2007) Medial prefrontal serotonin in the rat is involved in goal-directed behaviour when affect guides decision making. Psychopharmacology 195(3):435–449, DOI 10.1007/s00213-007-0917-6, URL https://doi.org/10.1007/s00213-007-0917-6

Ras T, van de Ven M, Patterson-Kane EG, Nelson K (2002) Rats’ preferences for corn versus wood-based bedding and nesting materials. Laboratory Animals 36(4):420–425, DOI 10.1258/002367702320389080, URL https://doi.org/10.1258/002367702320389080

Richman CL, Dember WN, Kim P (1986) Spontaneous alternation behavior in animals: A review. Current Psychological Research & Reviews 5(4):358–391, DOI 10.1007/bf02686603, URL https://doi.org/10.1007/bf02686603

Roder JK, Roder JC, Gerlai R (1996) Conspecific exploration in the t-maze: Abnormalities in s100 beta transgenic mice. Physiology & Behavior 60(1):31–36, DOI 10.1016/0031-9384(95)02247-3, URL https://doi.org/10.1016/0031-9384(95)02247-3

Rooijen JV, Metz J (1987) A preliminary experiment on t-maze choice tests. Applied Animal Behaviour Science 19(1-2):51–56, DOI 10.1016/0168-1591(87)90202-4, URL https://doi.org/10.1016/0168-1591(87)90202-4

Rudeck J, Vogl S, Banneke S, Schönfelder G, Lewejohann L (2020) Repeatability analysis improves the reliability of behavioral data. PLOS ONE 15(4):e0230900, DOI 10.1371/journal.pone.0230900, URL https://doi.org/10.1371/journal.pone.0230900

Sánchez-Santed F, de Bruin JP, Heinsbroek RP, Verwer RW (1997) Spatial delayed alternation of rats in a t-maze: effects of neurotoxic lesions of the medial prefrontal cortex and of t-maze rotations. Behavioural Brain Research 84(1-2):73–79, DOI 10.1016/s0166-4328(97)83327-x, URL https://doi.org/10.1016/s0166-4328(97)83327-x

Sharma S, Haselton J, Rakoczy S, Branshaw S, Brown-Borg HM (2010a) Spatial memory is enhanced in long-living ames dwarf mice and maintained following kainic acid induced neurodegeneration. Mechanisms of Ageing and Development 131(6):422–435, DOI 10.1016/j.mad.2010.06.004, URL https://doi.org/10.1016/j.mad.2010.06.004

Sharma S, Rakoczy S, Brown-Borg H (2010b) Assessment of spatial memory in mice. Life Sciences 87(17-18):521–536, DOI 10.1016/j.lfs.2010.09.004, URL https://doi.org/10.1016/j.lfs.2010.09.004

Shettleworth SJ (2010) Cognition, Evolution, and Behavior, 2nd edn. Oxford University Press, URL https://books.google.de/books?id=-Qs1qGys0AwC

Shoji H, Hagihara H, Takao K, Hattori S, Miyakawa T (2012) T-maze forced alternation and left-right discrimination tasks for assessing working and reference memory in mice. Journal of Visualized Experiments (60), DOI 10.3791/3300, URL https://doi.org/10.3791/3300

Sulikowski D, Burke D (2010) When a place is not a place: encoding of spatial information is dependent on reward type. Behaviour 147(11):1461–1479, DOI 10.1163/000579510x521564, URL https://doi.org/10.1163/000579510x521564

Tellegen A, Horn JM, Legrand RG (1969) Opportunity for aggression as a reinforcer in mice. Psychonomic Science 14(3):104–105, DOI 10.3758/bf03332727, URL https://doi.org/10.3758/bf03332727

Wadhera D, Wilkie LM, Capaldi-Phillips ED (2017) The rewarding effects of number and surface area of food in rats. Learning & Behavior 46(3):242–255, DOI 10.3758/s13420-017-0305-y, URL https://doi.org/10.3758/s13420-017-0305-y

Walker EL, Dember WN, Earl RW, Karoly AJ (1955) Choice alternation: I. stimulus vs. place vs. response. Journal of Comparative and Physiological Psychology 48(1):19–23, DOI 10.1037/h0047218, URL https://doi.org/10.1037/h0047218

Wenk GL (1998) Assessment of spatial memory using the t maze. Current Protocols in Neuroscience 4(1):8.5B.1–8.5A.7, DOI 10.1002/0471142301.ns0805bs04, URL https://doi.org/10.1002/0471142301.ns0805bs04

Wu CYC, Lerner FM e, Silva AC, Possoit HE, Hsieh TH, Neumann JT, Minagar A, Lin HW, Lee RHC (2018) Utilizing the modified t-maze to assess functional memory outcomes after cardiac arrest. Journal of Visualized Experiments (131), DOI 10.3791/56694, URL https://doi.org/10.3791/56694

Zhang Q, Kobayashi Y, Goto H, Itohara S (2018) An automated t-maze based apparatus and protocol for analyzing delay- and effort-based decision making in free moving rodents. Journal of Visualized Experiments (138), DOI 10.3791/57895, URL https://doi.org/10.3791/57895

