## Supplements for "Alternate without alternative: Neither preference nor simple learning behaviour shown by C57BL/6J mice in the T-maze"

#### 1 Transponder Implantation

At the age of five weeks, transponders (FDX-B transponder according to ISO 11784/85; group 1: Planet-ID, Germany; group 2: Euro I.D., Germany) were implanted under the skin in the neck of the mice. To do so, in group 1 all mice obtained an analgesic (Meloxicam) two hours before the procedure. The transponder implantation itself was performed under isoflurane anesthesia. RFID (radio frequency identification) transponders were injected directly behind the ears subcutaneously in the neck, so that they were rostrocaudal oriented. After transponder implantation, mice were placed in a separate cage with bedding and sheets of paper, and monitored until they were fully awake again. Then they were returned to their home cage. In group 1, two mice lost their transponders after the first implantation, and for those two mice the transponder implantation was repeated at the age of 8 weeks.

For group 2, the administration time of the analgesic was altered to the evening before the procedure because we hoped to reduce transponder loss this way: By administering the Meloxicam earlier, the analgesic effect was expected to cease before the dark phase after the implantation (active phase), and mice would be more hesitant to focus on the injection side. Implantation of the

Anne Habedank
German Federal Institute for Risk Assessment (BfR), German Center for the Protection of Laboratory Animals (Bf3R), Diedersdorfer Weg 1, D-12277 Berlin, Germany


Pia Kahnau
German Federal Institute for Risk Assessment (BfR), German Center for the Protection of Laboratory Animals (Bf3R), Diedersdorfer Weg 1, D-12277 Berlin, Germany

Lars Lewejohann
German Federal Institute for Risk Assessment (BfR), German Center for the Protection of Laboratory Animals (Bf3R), Diedersdorfer Weg 1, D-12277 Berlin, Germany
Institute of Animal Welfare, Animal Behavior and Laboratory Animal Science, Freie Universität Berlin, Königsweg 67, D-14163 Berlin, Germany

transponders was performed in the same way as in group 1. In group 2, no transponder was lost.

### 2 Experiment 1

#### 2.1 Tested goods

Two fluids ware compared, namely almond milk (3 g sugar per 100 ml, "Mandel drink", Alpro GmbH, Germany) and apple juice (100 %, 10 g sugar per 100 ml, made out of concentrate, Solevita, Lidl GmbH & Co. KG, Germany). We chose fluids because their odours can work as additional cue without previous conditioning.
During the eighth day of habituation, the two fluids were presented inside the home cage system: One of the usually two water bottles in each cage was replaced by a bottle containing one of the test fluids (500 ml). Originally, it was planned to present the bottles for a few days with randomised positions. However, after the first 24 h the bottle with the almond milk was nearly empty. As there was no leakage of the bottle, we have to assume that the mice drank all of the missing fluid. Health of the mice seemed to be unaffected but we noted excessive urination inside the home cages and the T-maze. Therefore, presentation of the two fluids was immediately stopped.

#### 2.2 Habituation to the T-Maze

Mice were habituated passively to the T-maze for 13 days, during which they could enter the T-maze whenever they were motivated. To do so, the tube between the two home cages was interrupted by a junction, which had a connection to the T-maze via a tube (40 mm diameter). RFID antennas were installed to receive information on the mice visiting the T-maze. Because mice were too fast for the RFID antennas, they were slowed down by two doors. After a first 15 cm long tube followed one door, then a 40 cm long tube, a second door, and a 6 cm long tube leading into the T-maze. Each door was directed by an Arduino micro-controller and surrounded by a light barrier (outer side, leading to the home cage or the maze) and an RFID antenna (inner side, leading to next door). Doors opened for 5 s when the transponder of a mouse was detected by the RFID antenna or a mouse interrupted the light barrier. In addition, with the help of two RFID readers also the direction of movement was reconstructable. In this manner, mice could move freely in and out of the T-maze, while their individual stay time was monitored via the RFID readers and stored onto an SD card by the Arduino.
Every day (except for the weekends), the maze was detached from the cage system, washed with water and then cleaned with 70 % ethanol to "reset" odour conditions. After the 13th day of habituation to the T-maze, mice cages were transported from their husbandry room to the experimental room. Here,

mice had one day to habituate to the new environment before the start of the experiment. Note that this was also already an extended habituation time as the common T-maze protocols recommend from 10 min up to over 30 min for habituation to the test room.

### 2.3 Preference Test

In experiment 1, the T-maze test (not habituation) took place in an experimental room, and an additional light was placed above the T-maze. Light conditions for the left arm were 171 lux, 201 lux for the right arm, 350 lux for the start/central arm, and 264 lux for the spot between the choice arms.
T-maze testing was conducted on two consecutive days. Mice were habituated to the test room before (see above) and performed five test trials per day, with an additional habituation trial beforehand on trial day 1. Test duration was approximately 40 min per mouse, lasting about 9 h per trials day for the whole group.
The order of tested mice was randomised for both trial days. In addition, presentation side of fluids was randomised for the mice so that for half of the mice almond milk was presented in the left arm and apple juice in the right and for half of the mice the other way round. For trial day one, presentation side of the fluids did not change between trials. On trial day two, two trials were performed with fluids presented in the same arm as the day before, while in trials three to five presentation sides were reversed to control for a potential side preference.
Before each mouse, 1 ml of the test fluids was administered on a cellulose sheet and stuck to the walls at the end of a choice arm as an odour stimulus. In addition, a fluid droplet of 20 $\mu$l was placed on the floor of the respective arm as a reward.
All trials were recorded with a video camera (Logitech C390e, Switzerland) and iSpy 64 (version 7.0.3.0). Before each mouse, maze and start cage were cleaned with 70 % ethanol and bedding in the start cage was replaced by new bedding. Then, the additional light was switched on, and the automated system controlling the door was started.
The first trial of the first day was the habituation trial: A mouse was taken out of the home cage by tube handling and placed into the start cage. The mouse had now 5 min to initiate a trial by going through the tube into the T-maze. If the mouse did not enter the T-maze during this time, it was lifted by the handling tube, allowing the mouse only to leave the tube into the tube leading to the maze by blocking the other tube entry. It was then waited until the mouse had entered both arms of the maze (crossed the virtual line with the whole body but not yet with its tail). After additional 30 s, the mouse was returned to the start cage with the help of the handling tube. Before the start of the next trial, the light and the automated system controlling the door were switched off; this prevented the mice from entering the maze, while the floor was cleaned with 70 % ethanol.

After drying the maze and replacing the droplet on the floor, light and automated system were turned on again and the mouse could re-enter the maze. The mouse had 3 min to do before it was guided by tube handling into the maze. During the test trials, it was waited until a mouse had entered one of the arms (crossed the virtual line with the whole body but not yet with its tail), before it was returned to the start cage with the handling tube. After the last trial, the mouse was returned to its home cage. In-between mice, the whole maze including the walls were cleaned with 70 % ethanol.

On day two, there was no habituation trial. In addition, a side switch of the fluids took place after trial two; therefore, in-between trials not only the floor but the complete maze was cleaned with 70 % ethanol. Also not only to the droplet on the floor but also the cellulose sheet at the end of each arm was renewed.

For video recording, a webcam (Logitech C390e, Switzerland) was mounted above the maze on a metal beam construction. The connected computer was placed near the T-maze in such a way that the experimenter could observe the mouse in the T-maze via the computer screen.

#### 2.4 Additional Notes on the Analysis

During passive T-maze habituation, the two Arduinos automatically saved all RFID detections and additional events (door opened/closed or light barrier interrupted) onto an SD card. Each record included a time stamp (hours, minutes and seconds since start of the Arduino, provided by a real-time clock), milliseconds passed since start of the Arduino, type of event, and the unique RFID transponder number. With the help of R studio (Version 1.1.383), the data sets recorded by the two Arduinos were then tagged with a number for each Arduino and merged. To analyse mouse visits to the T-maze, position changes were extracted (whenever a mouse was detected first by one reader and then by the other), excluding all additional events and RFID detection duplicates (if the same In experiment 2, the test took place in the husbandry room and no additional light was added. This led to illumination levels of 18 lux minimum at the end of both arms and 50 lux maximum at the start arm. In experiment 2, choice arms of the maze were covered with patterns.mouse was detected multiple times). Using the time stamps it could then be analysed how long each mouse stayed inside the maze.

In total, 143 trials were analysed, containing 13 habituation trials. Of the 130 preference test trials, five could not be assessed due to camera problems (camera recording stopped unnoticed for one mouse), leaving 125 trials.

Originally, it was planned also to take into account whether the mouse had taken the reward droplet on the floor or not. However, for apple juice this was not possible: Because of its transparency (in comparison to almond milk), the wet floor left behind looked too similar to the apple juice droplet itself. Therefore, we instead assessed whether the mouse spent some time ($> 1$ s) in which it its behaviour suggested licking or intensely sniffing the droplet.

### 3 Experiment 2

#### 3.1 Tested goods

In pre-tests we observed that millet seems to be a better working reward than almond milk: While mice showed no interest in almond milk when offered in a separate cage filled with home cage bedding, mice immediately fed on millet grains. In addition, after a few sessions of habituation, mice also fed on millet in an empty type III macrolon cage within one minute after entering the cage. We therefore expected mice to do so in the T-maze after the habituation trials as well.
As the aim of this test was mainly to establish a working protocol for the preference test, we decided against comparison of millet and another reward. Instead, we tested millet against "nothing". In this manner, the test design also resembled a simple learning test. To control for the visual (or exploratory) effect, we provided millet mixed with a specific bedding material in one arm and bedding material (without millet) in the other arm. As bedding material we used the same bedding material as in the home cage (Lignocel FS14, spruce / fir, 2.5-4 mm, JRS, J. Rettenmaier & Söhne GmbH + Co KG, Germany) as this was a definitely neutral (familiar) cue. Mice were already habituated to the millet in the course of other experiments (including the pre-tests).

#### 3.2 Habituation to the T-Maze

While in the last experiment, mice were habituated passively to the T-maze, in this experiment mice were manually habituated to the maze: Mice were placed individually into the maze setup for a short time period on five consecutive days.
This time, the T-maze was installed in the same room in which the mice were usually kept, so no transportation was necessary. In this husbandry room, no other groups of mice were kept during this experiment. Habituation trials were performed between 8:00 and 11:00 in the morning. After preparation of the setup, the filter top of the home cage system was removed and mice had 10 min to habituate to the illumination change.
For habituation to the maze, mice were taken individually and in a randomized order out of the cage and placed into a start cage, which contained only bedding material and was connected to the T-maze via a tube with an automated door (similar to the setup in the last experiment). Mice had already experiences with automated doors, thus, no habituation to the door was needed. Starting at the moment the mice entered the T-maze, they had 3 min to explore the whole maze. A return to the start cage was blocked by the automated door. If a mouse did not enter the maze within 3 min, it was retrieved by tube handling and held in front of the connection tube with the end to the start cage closed. If it then again did not enter the T-maze within the next 7 min, it was placed directly inside the maze. After 3 min of T-maze exploration, mice

were returned to their home cage.
During habituation, maze arms were empty and without visual cues. The maze was not disinfected in-between mice but it was cleaned (using paper and water) whenever defecation or urination were observed. The exploration behaviour inside the maze was recorded by a video camera (Logitech C390e, Switzerland) mounted above the maze on a metal beam construction.
It has to be noted that one mouse of the twelve received only four days of habituation: On day one it showed unusual behaviour which might have been correlated with health issues, and therefore, was excluded. As its behaviour returned to normal within two hours (and the maze test does not cause any severity), and the veterinarian had no objection, we decided to start habituation with this mouse on day two. In the course of the following three weeks (one habituation week and two test weeks) there was no unusual behaviour observed.

### 3.3 Preference Test

In experiment 2, the test took place in the husbandry room and no additional light was added. This led to illumination levels of 18 lux minimum at the end of both arms and 50 lux maximum at the start arm. In experiment 2, choice arms of the maze were covered with patterns.
While in the last experiment preference tests were conducted block-wise (two days with five trials each), in experiment 2 preference tests were conducted on five consecutive days, with only two trials (week 1) or three trials (week 2) per day. This test design should enable an improving habituation to the test with every experimental day. It also allowed flexible addition of trials if necessary (e.g., if mice had not fed on the millet due to still insufficient habituation).
Between habituation trials and test trials, there was a two day break. Just like the habituation, the preference test took place in the same room in which the mice were usually kept. Tests were performed between 8:00 and 11:00 in the morning, taking approximately 6 (week 1) to 8 (week 2) min per mouse. After preparation of the setup (installing laptop and cameras), the filter top of the home cage system was removed and mice had 10 min to habituate to the illumination change.
For the preference test trials, in one of the maze arms 0.05 g millet mixed with bedding material and in the other maze arm a similar amount of bedding material was placed. Walls of both arms were decorated with patterns: either white dots on black ground or white and black stripes. (Patterns are designed according to the description of Cunningham et al. 2006, except that the colour was inverted.) Combination of pattern, treatment and side were randomized across mice. In week 1, presentation side of the millet was kept the same for six trials, and then the side was switched (similar to experiment 1). In week 2, no side change was conducted.
Each experimental day, following a randomized order a mouse was taken out

of the cage and placed individually into a start cage. The start cage contained only bedding material and was connected to the T-maze via a tube with an automated door. The mouse now had 3 min to initiate a trial by entering the T-maze. If a mouse had not entered the maze within 3 min, it would have been retrieved by tube handling and held in front of the connection tube with the end to the start cage blocked. Entering the maze, the mouse had the choice between the rewarded (millet and bedding material) and the unrewarded arm (bedding material). As soon as the mouse crossed a virtual line which was 11 cm into the arm, this was considered a choice. The mouse was given time to feed on the millet, while leaving the arm was prevented by the experimenter's hand holding the handling tube. As soon as the mouse entered the tube, it was returned to the start cage and the procedure was repeated. After the second trial (or third trial, week 2) the mouse was returned to the home cage.
Between the two trials of the same mouse, the maze was not cleaned. Between different mice, the maze was not disinfected but it was cleaned (using paper and water) whenever defecation or urination were observed. Both trials were recorded by a video camera (Logitech C390e, Switzerland) mounted above the maze on a metal beam construction. As in experiment 1, the connected computer was placed near the T-maze in such a way that the experimenter could observe the mouse in the T-maze via the computer screen.

#### 3.4 Additional Notes on the Analysis

In total, 300 trials were analysed (10 per mouse in week 1, 15 in week 2). Of the 300 preference test trials, one missed the time point of the mouse entering the start cage because the video recording started too late.
Originally, it was planned to also take into account how long the mouse spent eating on the millet. However, as in all but two cases the millet was eaten completely (at least as far as visible) and mice had very different feeding speed, we decided against it.

### References

Cunningham CL, Gremel CM, Groblewski PA (2006) Drug-induced conditioned place preference and aversion in mice. Nature Protocols 1(4):1662–1670, DOI 10.1038/nprot.2006.279, URL https://doi.org/10.1038/nprot.2006.279
